# Mechanistic insight into substrate specificity of plant glucosinolate transporters

**DOI:** 10.1101/2023.01.06.522984

**Authors:** Christa Kanstrup, Nikolai Wulff, Carlos Peña-Varas, Morten Egevang Jørgensen, Rose Bang-Sørensen, Christoph Crocoll, Flemming Steen Jørgensen, David Ramírez, Ingo Dreyer, Osman Mirza, Hussam H. Nour-Eldin

## Abstract

Plants depend on transport processes for correct allocation of specialized metabolites. This is important for optimal defense, avoidance of autotoxicity, connecting compartmented biosynthetic modules and more. Transport of a wide variety of specialized metabolites is mediated by transporters from the Nitrate and Peptide transporter Family (NPF), which belongs to the Major Facilitator Superfamily (MFS). However, the mechanism by which NPF members recognize and transport specialized metabolites remains unknown.

Here we mutate eight residues to reciprocally swap the substrate-preference of two closely related glucosinolate transporters (GTRs). Seven of these residues assemble in a ring-like structure in all conformations of the transporters. We labeled the ring-like structure a selectivity filter and based on docking studies, we propose that the interaction between the selectivity filter and the glucosinolate side chain determines whether a given glucosinolate is recognized as a substrate. Besides partly explaining the distinct substrate preference of GTR1 (NPF2.10) and GTR3 (NPF2.9), this study proposes fundamental principles of substrate recognition in the NPF and establishes the GTR subclade as a novel model system for studying structure function relationships in the NPF.

## Introduction

In plants, specialized metabolism is estimated to produce between 200.000 to 1 million different chemical structures that enable plants to interact with their ever-changing environment and to overcome both biotic and abiotic stresses (Fang et al. 2019, Wang et al. 2019). Accordingly, the intricate involvement of transport processes in synthesis, storage, mobilization and conversion of specialized metabolites has necessitated the evolution of transporters capable of transporting an immense diversity of chemical structures.

The Nitrate and Peptide transporter Family (NPF) is a subfamily of the proton dependent oligopeptide transporter family (POT) (Léran et al. 2014), whose few members in humans are essential for uptake of di- and tripeptides with nearly limitless variations in amino acid side chains resulting from dietary digestion of proteins (Martinez Molledo et al. 2018). Based on extensive structural investigations of bacterial and mammalian homologs, this promiscuity can be attributed to key features, namely, a few conserved that interact with conserved features of the di- and tripeptide backbones and a large substrate-binding cavity that accommodate the rest of the oligopeptide residues (Doki et al. 2013, Guettou et al. 2013, 2014; Killer et al. 2021, Lyons et al. 2014, Parker et al. 2021).

In plants, the NPF family has undergone a tremendous expansion in terms of gene number. For example, in *Arabidopsis*, rice and tomato, the NPF family comprises 53, 70 and 77 members, respectively (Longo et al. 2018). The family consists of eight clades that are conserved in all plants. Over the past decade, NPF clades 1 and 2 have emerged as hotspots for transporters of almost all classes of specialized metabolites. As of now the list of substrate classes include, glucosinolates, cyanogenic glucosides, steroid glycoalkaloids, monoterpene indole alkaloids and flavonoids (Kanstrup & Nour-Eldin 2022). Hence, a multitude of physiological roles that so far include delivery of defense compounds to seeds (Nour-Eldin et al. 2012, 2017), detoxification of maturing fruits (Kazachkova et al. 2021) and linking compartmentalized biosynthetic modules (Payne et al. 2017) reflects a remarkable ability of NPF-clades one and two to evolve novel substrate specificities. In plants, resolving the atomic structure of the founding member the nitrate transporter NPF6.3 (Parker & Newstead 2014, Sun et al. 2014), pinpointed residues involved in binding nitrate but did not reveal how specialized metabolites are recognized by NPF members.

Glucosinolates are a class of *Brassica* specific amino acid-derived defense compounds. The ~130 known glucosinolates are categorized by the chemical structure of the parent amino acid. Those derived from methionine and tryptophan are categorized as aliphatic and indole glucosinolates, respectively, and those derived from phenylalanine and tyrosine as aromatic glucosinolates (Halkier & Gershenzon 2006). The glucosinolate transporters AtNPF2.10 and AtNPF2.11 from *A. thaliana* share ~80 % amino acid sequence identity and transport all glucosinolates regardless of parent amino acid side-chain (Andersen et al. 2013, Jørgensen et al. 2017, Nour-Eldin et al. 2012). In contrast AtNPF2.9, which is ~60 % identical to NPF2.10/NPF2.11, displays a strong preference for indole glucosinolates (Jørgensen et al. 2017). The NPF2.10/NPF2.11 and NPF2.9 anchored subclades are conserved across *Brassica* species. We hypothesize that the core part of the structurally diverse glucosinolates, i.e. the thioglucose and sulfate moieties, interact with the same residues in NPF2.10, NPF2.11 and NPF2.9 transporters and that the difference in substrate preference is due to differences in residues interacting with the varying side-chains. Hence, we reasoned that we could identify the residues that are involved in binding glucosinolates in the NPF2.9-2.11 by pursuing a mutational gain-of-function approach that converts the narrow substrate selectivity of NPF2.9 towards the broader preference of NPF2.10 and vice versa.

Here, we show experimentally that the substrate preferences of NPF2.10 and NPF2.9 can be swapped by mutating 8 residues within the substrate binding cavity and that seven of these residues form a ring-like selectivity filter in the substrate binding cavity. Docking studies support that interaction between the selectivity filter and glucosinolate side chain determines whether a given glucosinolate is a substrate and pinpoints the residues interacting with the glucosinolate core parts. This study establishes the glucosinolate transporters as a novel model system for unraveling the molecular basis for substrate recognition by the NPF family. Our findings lead us to propose a mechanistic model for how NPF members discriminate between substrates belonging to plant specialized metabolism.

## Results

### Differentially conserved residues in glucosinolate transporters

As a first step, we identified orthologues for each of the three *A. thaliana* glucosinolate transporters NPF2.10, NPF2.11 and NPF2.9 in other glucosinolate producing plants. From *A. thaliana, Brassica napus, Brassica rapa, Thellungiella halophila, Eutrema halophilum, Arabidopsis lyrata, Boechera stricta, Capsella rubella* and *Capsella grandiflora*, 13 non-redundant orthologues were selected for each glucosinolate transporter by using their respective protein sequence as BLAST query on the NCBI (Johnson et al. 2008) and Phytozome (Goodstein et al. 2012) servers. The phylogenetic tree of the 39 sequences was used to run a Type II Divergence test (Gu & Vander Velden 2002), which identified 72 amino acid positions that represented conserved differences comparing the 13 NPF2.9 sequences to the 26 NPF2.10/NPF2.11 sequences (Supplementary Table 1).

As a second step, we investigated whether the distinct substrate preference of the *A. thaliana* glucosinolate transporters extended across each subclade. We tested the substrate preference of a distant and a close member of the NPF2.10 and NPF2.9 subclade (Brara_A02314 and Carubv10016825M for NPF2.10, and Brara_F01308 and Carubv10012216M for NPF2.9) by expressing each orthologue in *Xenopus laevis* oocytes and exposing to either 100 μM 4-methylthio-3-butenyl glucosinolate (4MTB - aliphatic glucosinolate) or 100 μM indole-3-methyl glucosinolate (I3M - indole glucosinolate) for 1 hour at pH 5 (Fig. 1). The relative amounts of imported glucosinolates showed that the preference for I3M extended to the tested members of the NPF2.9 subclade and that the non-discriminating preference was preserved in the tested members of the NPF2.10 subclade.

**Fig. 1.**
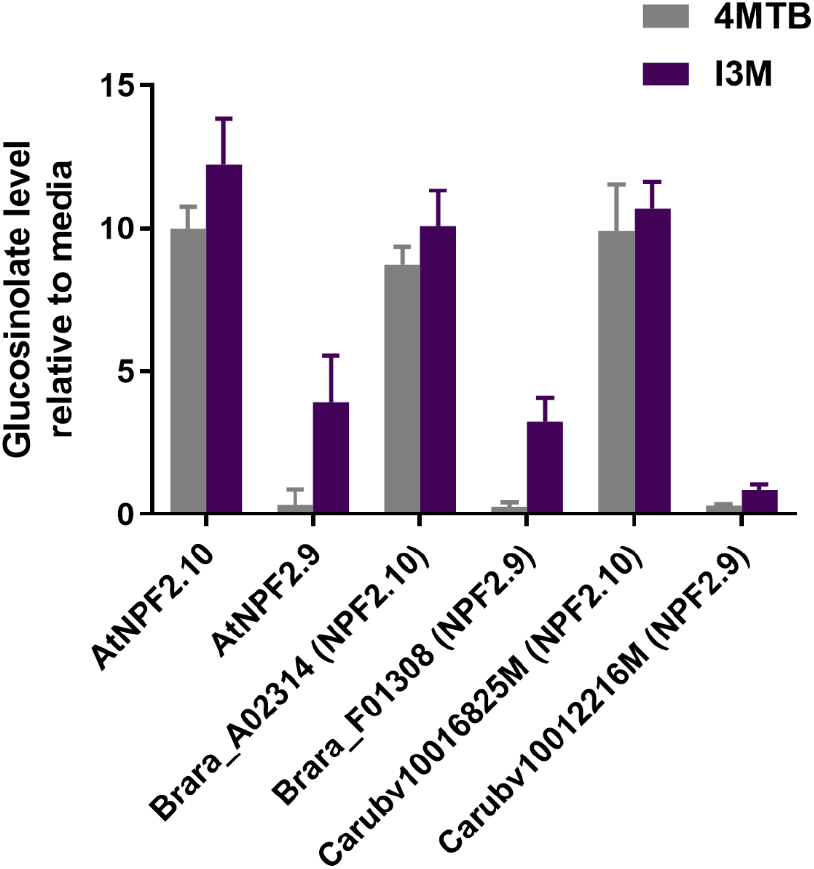
4MTB and I3M uptake of NPF2.9 and NPF2.10 orthologues. *Xenopus* oocytes expressing AtNPF2.10, AtNPF2.9, Brara_A02314, Brara_F01308, Carubv10016825M or Carubv10012216M were exposed to 100 μM 4MTB or 100 μM I3M for 60 min in pH 5 and glucosinolate content was quantified on LCMS (n = 4-6). The data is represented as relative to the external concentration of the glucosinolates, measured on LCMS.

### Differentially conserved residues relevant for substrate binding

We hypothesize that the distinct substrate preference displayed by NPF2.9 compared to NPF2.10/NPF2.11 can be explained by a subset of conserved differences within the 72 residues between the two subclades. To test this hypothesis, we aimed to convert the narrow I3M preference of NPF2.9 to the broad non-discriminating preference of NPF2.10 and vice versa. If all possible combinations of mutations within the 72 residues that are differently conserved between NPF2.9 and NPF2.10/NPF2.11 should be tested individually and in combination, it would require 4.72*10^21^ (2^72^) different mutants.

Recently, we pinpointed 51 residue positions that line the central substrate binding cavity of NPF proteins (Wulff et al. 2019). It is likely that surface exposed residues within the substrate binding cavity of the NPF transporters play an important role in substrate recognition as observed for transporters in the POT family (Doki et al. 2013, Guettou et al. 2013, 2014; Killer et al. 2021, Lyons et al. 2014, Parker et al. 2017, 2021). Hence, we focused on residues lining the substrate binding cavity and that are conserved differently between the NPF2.10/NPF211 and NPF2.9 clades. 16 of the 72 differently conserved residues are among the 51 residue positions in the central cavity (Fig. 2, Supplementary Table S1). Using a complementary approach for identifying potential substrate binding residues, we subjected homology models of NPF2.9 and NPF2.10 in the inward facing conformations (Jørgensen et al. 2017) to a HOLE analysis (Smart et al. 1996) and docked 24 structurally different glucosinolates (Supplementary Fig. S1) in the cavity using the Glide docking program (Friesner et al. 2006). These analyses identified 99 unique residue positions of which 16 are part of the 72 differently conserved residues (Fig. 2, Supplementary Table S1). 11 of these residues represent a common denominator between both analyses and were selected for mutational studies. In addition to these 11 common denominator residues, position 70 (in NPF2.9) was included as a control for sample handling (see Materials and Methods). The 12 positions are displayed in Fig. 3.

**Fig. 2.**
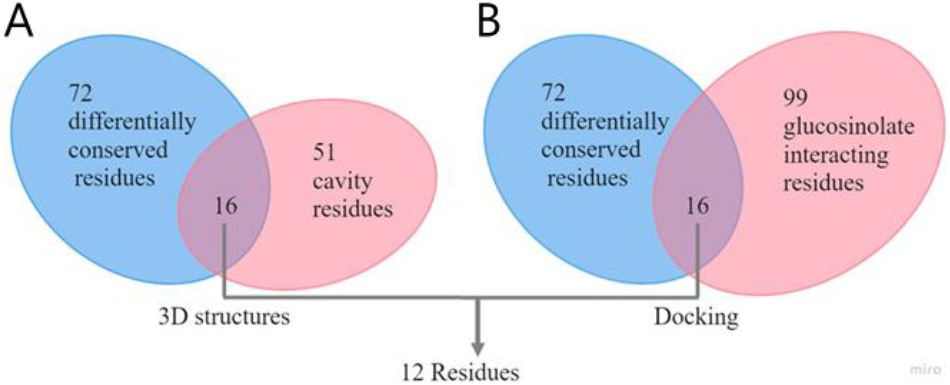
Identification of differentially conserved residues potentially responsible for the GTR3 and GTR1/GTR2 selectivity derived from the structure-based analysis (A) and by docking (B). Analysis A and B revealed 11 common residues selected for mutational studies. In addition position 70 in AtGTR3 (identified in both analysis A and B, but nor differently conserved) was included as a control for sample handling (see Materials and Methods).

**Fig. 3.**
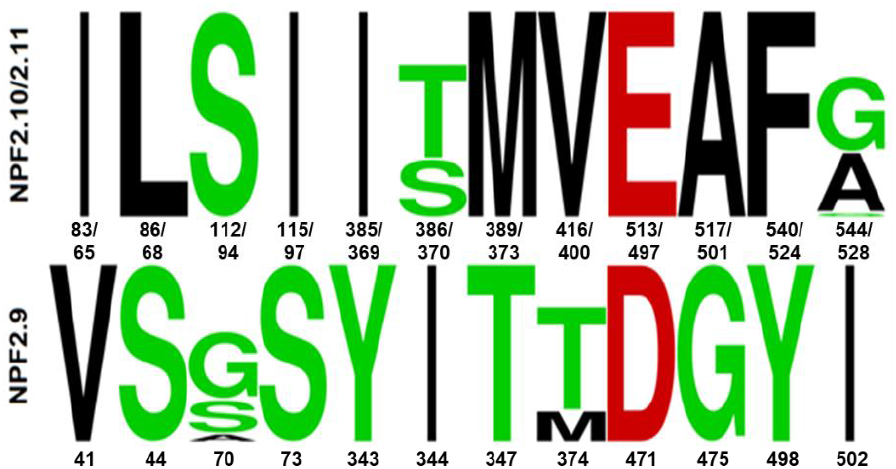
The identity of the 12 residue positions in the 39 glucosinolate transporter sequences presented as Logos. The top LOGO was created from the 26 GTR1/2 sequences from *A. thaliana, Brassica napus, Brassica rapa, Thellungiella halophila, Eutrema halophilum, Arabidopsis lyrata, Boechera stricta, Capsella rubella* and *Capsella grandiflora*. The numbers correspond to positions in AtGTR1/AtGTR2 (NPF2.10/2.11). The bottom LOGO from 13 GTR3 sequences from the same plants. The numbers correspond to positions in AtGTR3 (NPF2.9). The Logos were produced with the WebLogo server (Crooks et al. 2004)

### Narrowing the subset of residues involved in glucosinolate substrate specificity of NPF2.9

To assess whether the 72 and 12 residue-subsets are involved in determining substrate specificity, we introduced the NPF2.10 versions of the 12 and the full set of 72 residues, respectively, into *NPF2.9* sequence. The two NPF2.9 mutant versions were named NPF2.9Δ12 and NPF2.9Δ72, respectively, and were expressed in *Xenopus laevis* oocytes and exposed to either 100 μM aliphatic 4MTB or indole I3M for 60 min. NPF2.10 imported 4MTB and I3M into oocytes to similar levels (~4-5 fold above media level), whereas NPF2.9 imported I3M to ~20 fold higher levels than 4MTB. In comparison, the two mutant versions of NPF2.9 accumulated 4MTB and I3M in oocytes to equal levels indicating that we had converted the narrow substrate selectivity of NPF2.9 towards that of NPF2.10. However, the overall transport activity of both NPF2.9Δ12 and NPF2.9Δ72 were reduced by ~80 – 90 % compared to NPF2.9 (Fig. 4). These results indicated that the 12 mutations indeed affected substrate specificity but also had effects that impaired overall transport activity.

**Fig. 4.**
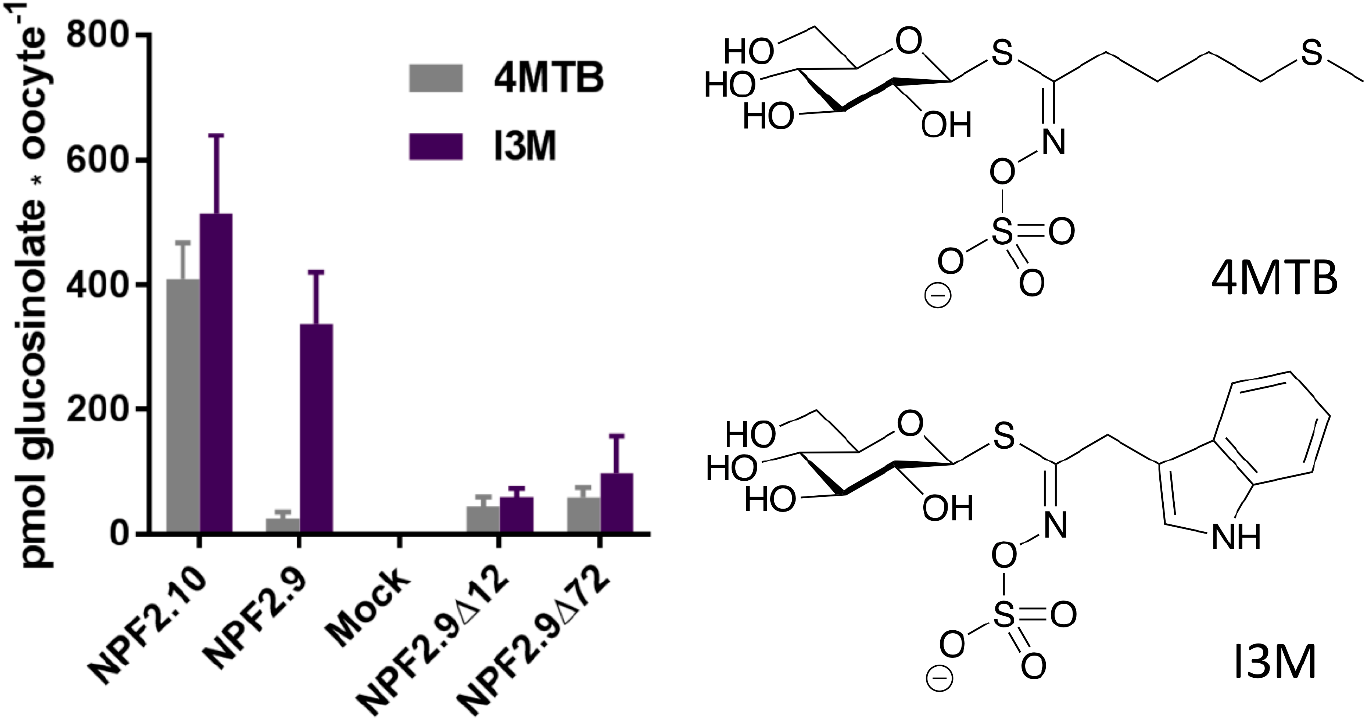
4MTB and I3M uptake of NPF2.9Δ12 and NPF2.9Δ72. *Xenopus* oocytes expressing NPF2.10, NPF2.9, NPF2.9Δ12, NPF2.9Δ72 or water-injected for control, were exposed to 100 μM 4MTB or 100 μM I3M for 60 min in pH 5 and glucosinolate content was quantified on LCMS (n = 5-6). The structures of 4MTB and I3M are displayed next to the bar diagram.

To identify the residues reducing transport rate we reverted each of the 12 residue mutations of NPF2.9Δ12 one-by-one to their original identity in NPF2.9; thus creating NPF2.9Δ11#1-#12 (Supplementary Table S2). These mutants are described using the following nomenclature: amino acid residue in NPF2.9 followed by the position in NPF2.9 and the version of the amino acid residue in NPF2.10. The version of the residue in the tested mutant is stated in bold.

All mutants were expressed in *Xenopus* oocytes and exposed to either 200 μM 4MTB or I3M for 60 min in pH 5. Three mutants, NPF2.9Δ11#5_**Y**343I, NPF2.9Δ11#7_**T**347M and NPF2.9Δ11#12_**I**502A, displayed increased activity compared to NPF2.9Δ12 (Fig. 5A) without affecting the gained NPF2.10-like non-discriminating preference. To explore potential additive effects we reverted the three residues in pairs (three combinations) and all three residues together. We exposed *Xenopus* oocytes expressing these four mutants to either 200 μM 4MTB or I3M for 60 min in pH 5. Both NPF2.9Δ10#_**T**347M_**I**502A and NPF2.9Δ9#1_**Y**343I_**T**347M_**I**502A-expressing oocytes were able to accumulate both 4MTB and I3M >3 fold over the media level. In comparison, NPF2.9 expressing oocytes only accumulated 4MTB to the media level and I3M to >4 fold over media level. Oocytes expressing NPF2.10 accumulated both 4MTB and I3M >6 fold over media level (Fig. 5B). Thus, by mutating only 9-10 residues that are conserved differently in NPF2.9 orthologues compared to NPF2.10/NPF2.11, we succeeded in changing NPF2.9 towards a broad specific glucosinolate transporter like NPF2.10/NPF2.11.

**Fig. 5.**
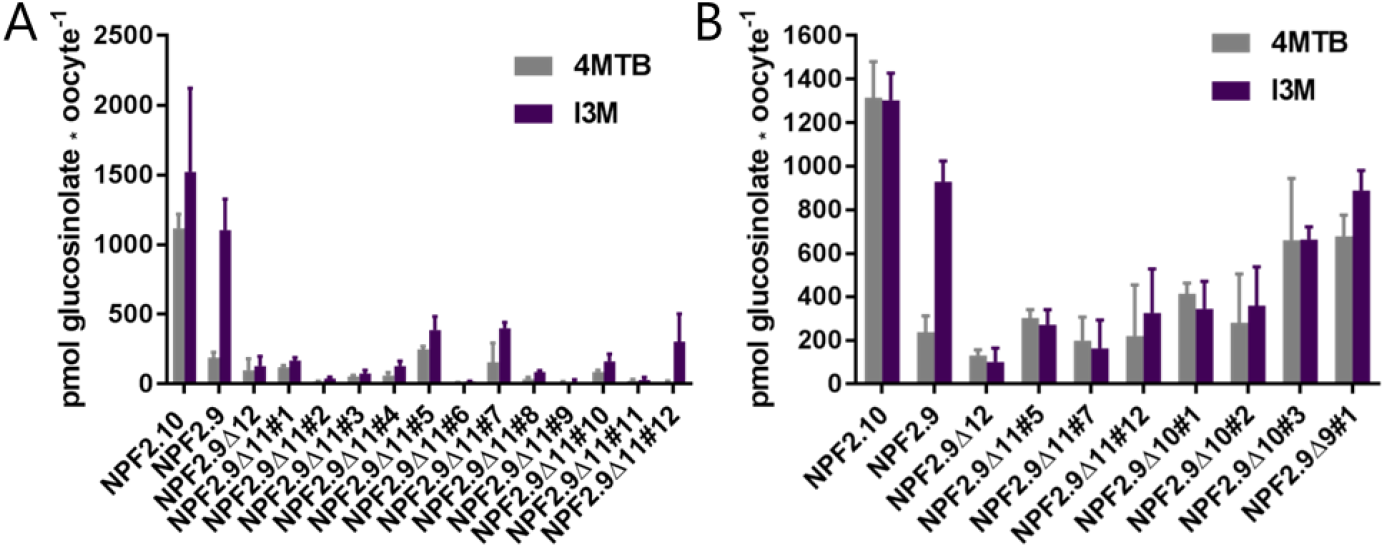
Activity of mutants. (**A**) Assessing the activity of NPF2.9 versions with 11 mutations; NPF2.9Δ11#1 to #12 is reverted in either of the 12 positions from NPF2.9Δ12 (n = 4-5). (**B**) Assessing the activity of NPF2.9 versions with 10 or 9 mutations. In combinations of hits from (**A**); NPF2.9Δ11#5 #7 and #12 (n = 5-6). Genes were expressed in *Xenopus* oocytes and exposing to 200 μM 4MTB or I3M for 60 min in pH 5. Glucosinolate content was quantified on LCMS.

### Equimolar competition assays reveals overall shift in substrate preference in the NPF2.9Δ10#3 mutant

So far the increased ability of the NPF2.9-to-NPF2.10 mutants to transport 4MTB was inferred from transport assays where 4MTB and I3M were used individually. To investigate substrate preference in higher detail we exposed NPF2.10, NPF2.9 and NPF2.9Δ10#3-mutant expressing oocytes to an equimolar mixture containing 13 different glucosinolates (Fig. 6). We chose to focus on investigating the substrate preference of the NPF2.9Δ10#3-mutant, because the slightly higher relative I3M uptake vs 4MTB (Fig 5B), indicated a more NPF2.9-like preference of the NPF2.9Δ9#1 mutant.

**Fig. 6.**
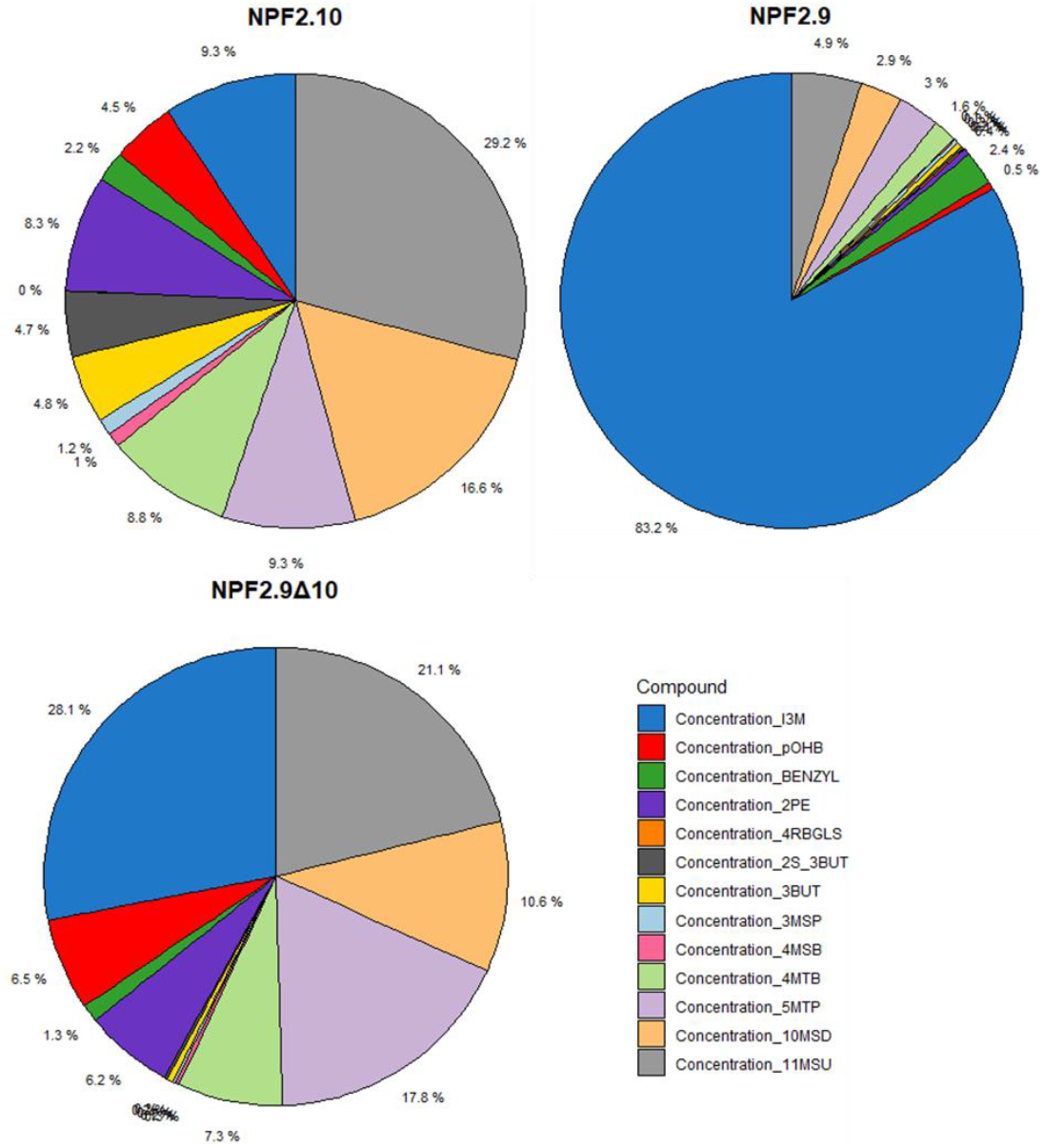
Equimolar competition assays of NPF2.9Δ10#3 with 13 glucosinolates. *Xenopus* oocytes expressing NPF2.10, NPF2.9, NPF2.9Δ10#3. Exposed to 100 μM I3M, pOHB, Benzyl, 2PE, 4RBGLS, 2S_3BUT, 3BUT, 3MSP, 4MSB, 4MTB, 5MTP, 10MSD, 11MSU in equimolar assay for 60 min pH 5. Glucosinolate content quantified on LCMS. Represented as Percent of each glucosinolate out of total in that oocyte. 1 oocyte batch, 5-6 oocytes per pie chart, analyzed as single oocytes.

The mixture of 13 glucosinolates contains, I3M and 4MTB and in addition also the tyrosine-derived, parahydroxybenzyl glucosinolate (pOHB), the phenylalanine-derived benzyl and 2-phenylethyl glucosinolates (Benzyl and 2PE), and a range of methionine-derived short-chain and long-chain glucosinolates including both thio, sulfinyl and unsaturated side chains (Supplementary Fig S2). If NPF2.10 transported all glucosinolates with similar preference irrespective of the glucosinolate side chain, each glucosinolate should represent approximately 7.7% of the imported glucosinolates. This was roughly seen for seven of the included glucosinolates. In comparison, the two long-chain 10-methylsulfinyldecyl (10-MSD) and 11-methylsulfinylundecyl glucosinolates (11-MSU) each represented 16.6% and 29.2% of all glucosinolates imported, whereas benzyl, the short chain 3-methylsulfinylpropyl (3-MSP) and 4-methylsulfinylbutyl glucosinolates (4-MSB) each represented 2.2%, 1.2% and 1% of all glucosinolates imported, respectively. Uptake of 4-(alpha-L-Rhamnosyloxy)benzyl glucosinolate (4RBGLS) was not detected. In comparison, the strong preference for I3M by NPF2.9 was further cemented with I3M representing 83.2% of all imported glucosinolates in NPF2.9-expressing oocytes. Analysis of the glucosinolates imported by the NPF2.9Δ10#3 mutant revealed a marked change in the substrate preference of NPF2.9. For example, I3M only represented 28.1% of all imported glucosinolates whereas 11-MSU represented 21.1% of all imported glucosinolates. Similar shifts toward NPF2.10-like preference patterns was seen for eight of the included glucosinolates.

### Identifying the minimal subset of residues required to broaden the substrate preference in NPF2.9

Next we hypothesized that the 10 mutated residues in the NPF2.9Δ10#3-mutant might include residues that neither influence the substrate preference nor the transport rate. Hence, to identify the minimal set of mutations necessary to broaden the substrate preference of NPF2.9 towards that of NPF2.10/NPF2.11 without affecting overall transport activity, we undertook another round of single residue reversions and screened for mutations that reversed the preference of the NPF2.9Δ10#3 mutant towards the preference of NPF2.9. We reverted the 10 mutations of NPF2.9Δ10#3 one-by-one by site-directed mutagenesis; creating NPF2.9Δ9#1-#10 (Supplementary Table S2). *Xenopus* oocytes expressing these 10 mutants were exposed to 200 μM pOHB for 60 min in pH 5 (Fig. 7). We switched to using pOHB because this glucosinolate - unlike 4MTB and I3M - is readily available at lower cost from commercial vendors. Three mutants NPF2.9Δ9#7_**V**374T, NPF2.9Δ9#8_**E**471D and NPF2.9Δ9#10_**F**498Y displayed NPF2.10-like uptake levels for pOHB, whereas (unwanted) NPF2.9-like uptake levels (i.e. reduced pOHB uptake) were seen in the rest of the mutant versions. Hence, the conversion of the T374, D471 and Y498 residues to their corresponding NPF2.10 version is not required for the broadening of NPF2.9 substrate preference (Fig. 7).

**Fig. 7.**
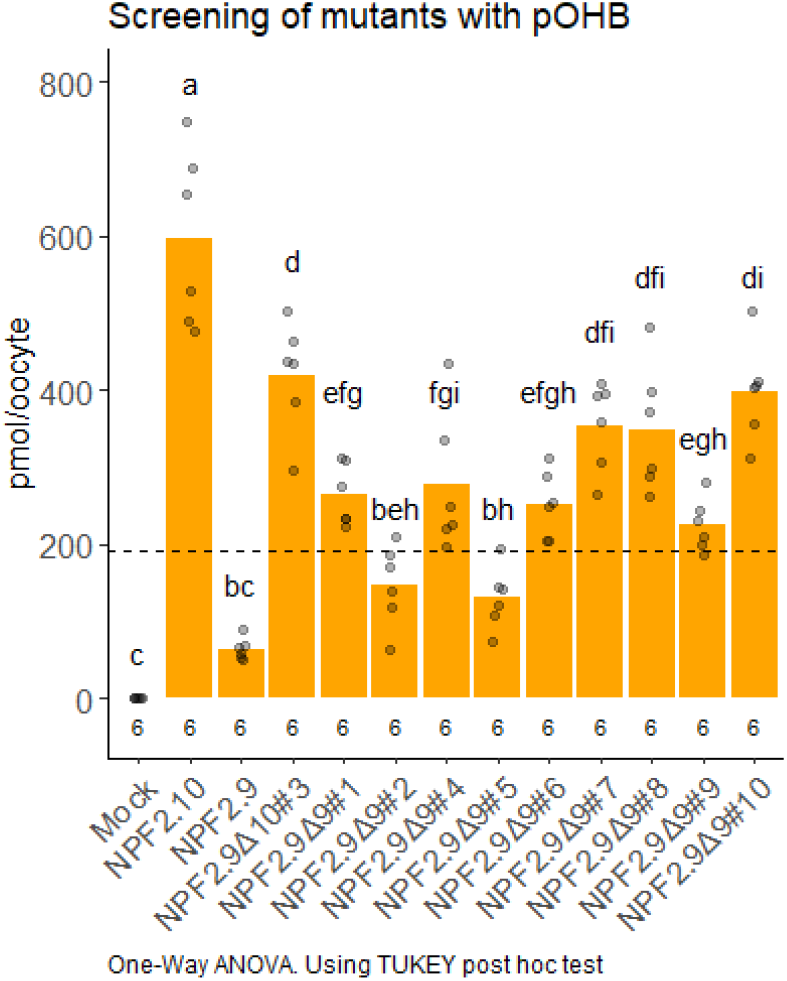
pOHB screen of NPF2.9Δ10#3 and NPF2.9Δ9 mutants. *Xenopus* oocytes expressing NPF2.10, NPF2.9, NPF2.9Δ10#3, 9 × NPF2.9Δ9 or water-injected mock for control. Exposed to 200 μM pOHB for 60 min pH 5. Glucosinolate content quantified on LCMS. Numbers in the bottom indicates number of oocytes over 1 batch, analyzed in single oocytes samples. Dotted lines represent the average media sample concentration. Statistical analysis: one-way ANOVA using TUKEY post hoc test comparing the transporters. Same letters indicate no significant difference

To explore whether the residues T374, D471 and Y498 could be omitted in combination, we reverted the three residues in pairs (three combinations) and all three residues together; NPF2.9Δ8#1_**V**374T_**E**471D, NPF2.9Δ8#2_**V**374T_**F**498Y, NPF2.9Δ8#3_**E**471D_**F**498Y and NPF2.9Δ7#1_**V**374T_**E**471D_**F**498Y (Supplementary table S2). *Xenopus* oocytes expressing the different NPF2.9Δ8-mutants and the NPF2.9Δ7 mutant were exposed to pOHB alone and a mixture of 200 μM 4MTB, I3M and pOHB, respectively, for 60 min in pH 5 (Fig. 8). In the single pOHB assay, both NPF2.9Δ8#2, NPF2.9Δ8#3 and NPF2.9Δ7#1 expressing oocytes accumulated pOHB to similar levels as the NPF2.9Δ10#3; ~2-fold over media level. NPF2.9 expressing oocytes accumulated pOHB below media level (Fig. 8A). When exposed to the mixture of the three glucosinolates, both NPF2.9Δ8#2, NPF2.9Δ8#3 expressing oocytes were able to accumulate I3M above media level, and pOHB to around media level, similar to the accumulation pattern of NPF2.10. In comparison, the accumulation of 4MTB by the mutants reached intermediate levels compared to NPF2.10 (Fig. 8B). Surprisingly, the NPF2.9Δ8#1_**V**374T_**E**471D, lowered the accumulation of pOHB to below media level in the assay where pOHB was included alone (Fig. 8A), and lowered the overall uptake of all glucosinolates when in combination (Fig. 8B). With the slightly lowered uptake of all three glucosinolates by the NPF2.9Δ7#1 mutant it might suggest the combination of V374 & E471 negatively affects the specificity and transport rate.

**Fig. 8.**
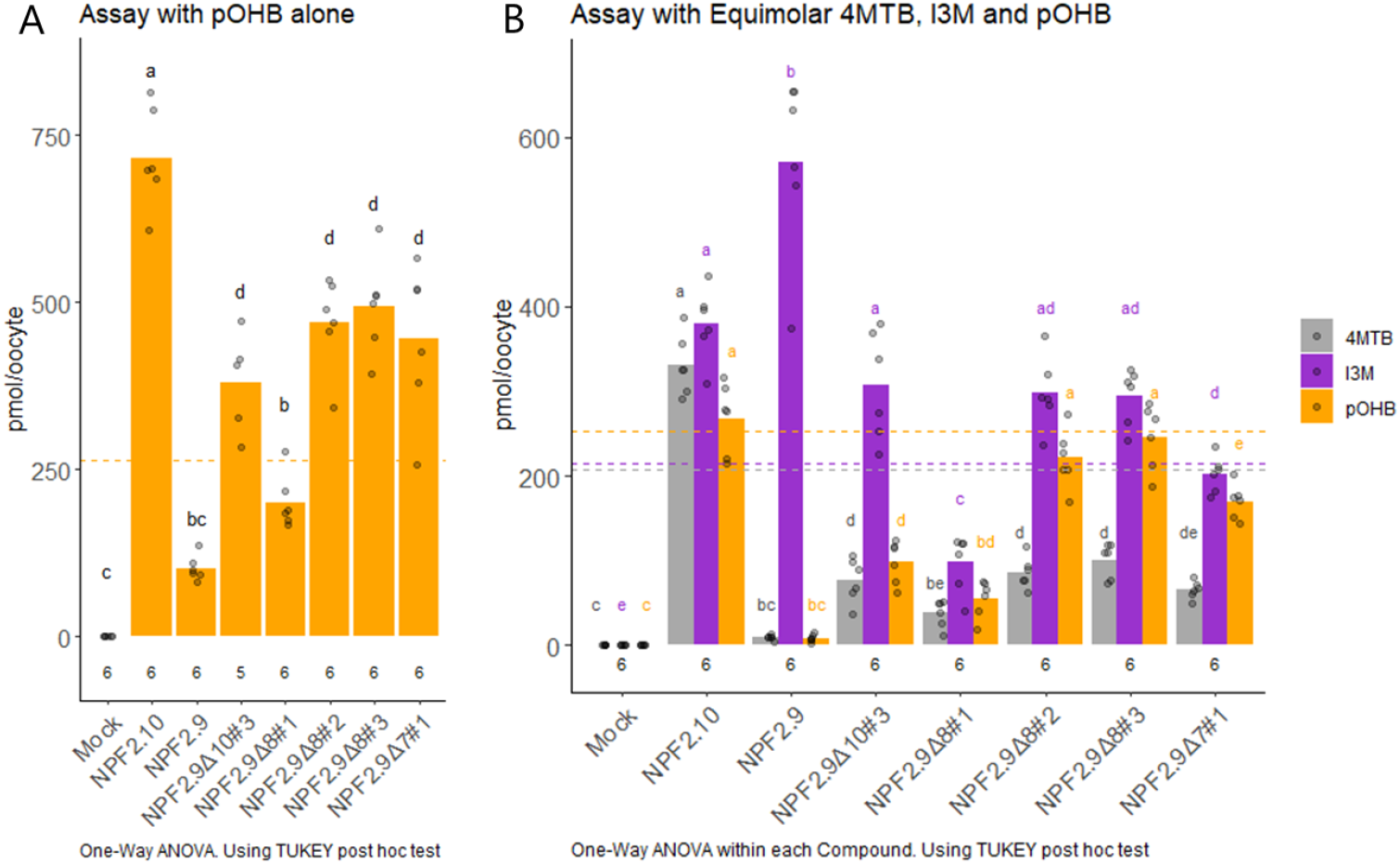
4MTB, I3M and pOHB uptake of NPF2.9Δ8 and NPF2.9Δ7. *Xenopus* oocytes expressing NPF2.10, NPF2.9, NPF2.9Δ10#3, 3 × NPF2.9Δ8, NPF2.9Δ7 or water-injected mock. A) Oocytes exposed to 200 μM pOHB for 60 min pH 5. B) Oocytes exposed to 200 μM 4MTB, I3M and pOHB in equimolar assay for 60 min pH 5. Glucosinolate content quantified on LCMS. Numbers in the bottom indicates number of oocytes over 1 batch, analyzed in single oocytes samples. Dotted lines represent the average media sample concentration. Statistical analysis: one-way ANOVA for each compound using TUKEY post hoc test. In A, all transporters compared. In B transporters compared for each compound, color on labels indicate compound. Same letters indicate no significant difference, only same color should be compared (within compound).

These transport assays revealed that both the NPF2.9Δ8#2 and NPF2.9Δ8#3 mutants displayed NPF2.10-like substrate preferences towards pOHB and I3M whereas 4MTB uptake was increased to an intermediate level, and not significantly higher than the NPF2.9Δ10#3. Thus, these data illustrate that the eight mutations in the NPF2.9Δ8#2 and NPF2.9Δ8#3 mutants largely convert the substrate preference of NPF2.9 into that of NPF2.10/NPF2.11.

### Mutating the equivalent residues in NPF2.10 change the substrate specificity for pOHB, but not for 4MTB

The introduction of the eight mutations in NPF2.9, broadens the substrate specificity of NPF2.9 to match that of NPF2.10/NPF2.11 by allowing transport of pOHB to almost the same extent as I3M. This prompted us to investigate whether introducing the reciprocal mutations in NPF2.10 would reduce the preference of NPF2.10 towards pOHB. We chose the residues in the NPF2.9Δ8#3-mutant for introducing the reciprocal mutations in NPF2.10, due to the slightly higher (though not significantly different) uptake of 4MTB and pOHB (Fig. 8AB). The eight mutations were introduced in NPF2.10 (NPF2.10Δ8) (Supplementary table S3) and NPF2.10Δ8 was expressed in *Xenopus* oocytes side by side with NPF2.10 and NPF2.9. Expressing oocytes were exposed to mixture of 200 μM 4MTB, I3M and pOHB for 60 min in pH 5 (Fig. 9). NPF2.10Δ8 expressing oocytes were able to accumulate both 4MTB and I3M to the same extent as NPF2.10 - above media level. In comparison, the pOHB uptake is almost completely abolished like for NPF2.9.

**Fig. 9.**
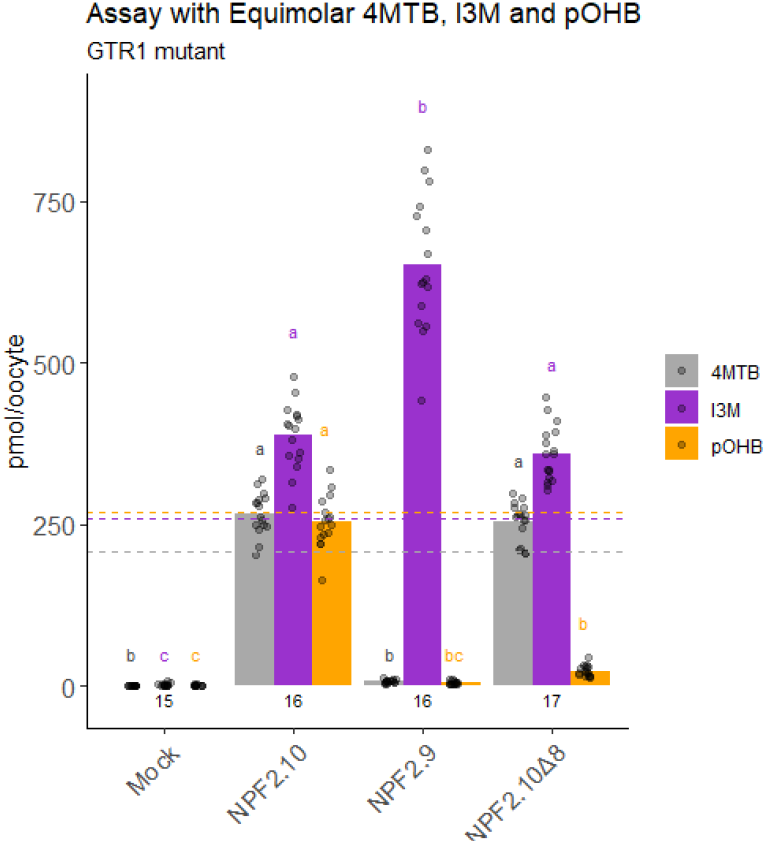
4MTB, I3M and pOHB uptake of NPF2.10Δ8. *Xenopus* oocytes expressing NPF2.10, NPF2.9, NPF2.10Δ8 or water-injected mock exposed to 200 μM 4MTB, I3M and pOHB in equimolar assay for 60 min pH 5. Glucosinolate content quantified on LCMS. Numbers in the bottom indicates number of oocytes over 3 batches, analyzed as single oocytes. Dotted lines represent the average media sample concentration. Statistical analysis: one-way ANOVA for each compound using TUKEY post hoc test comparing the transporters. Same letters indicate no significant difference, only same color should be compared (within compound).

Introducing the eight mutations into NPF2.10 does not change the substrate preference for 4MTB and I3M, however, the mutant shows a complete NPF2.9-like preference for pOHB. Thus, these data illustrate that the eight mutations in the NPF2.10Δ8 mutant converts the substrate preference of NPF2.10 into that of NPF2.9 for pOHB.

### Generation of homology models for structural interpretation of mutational data

In order to facilitate a structural interpretation of the eight mutations identified in NPF2.9Δ8#3, we docked I3M and pOHB in NPF2.9, NPF2.9Δ8#3 and NPF2.10 homology models. We have previously performed dockings of 4MTB and I3M in outward and inward facing conformations of NPF2.10 (Jørgensen et al. 2017, Peña-Varas et al. 2022). In this study, we were interested in performing the dockings in an occluded state, to better study the binding cavity. We browsed through Alpha Fold models (Jumper et al. 2021, Varadi et al. 2022) of *A. thaliana* NPFs, and found 1 predicted model to be close to an occluded state: AtNPF4.4 (Identifier AF-Q56XQ6-F1). This model was chosen as a template for homology modeling of NPF2.9, NPF2.9Δ8#3 and NPF2.10. The models were created and validated using SWISS-MODEL. After generation of the models, they were relaxed through Molecular Dynamics simulations (MDs) as previously described (Peña-Varas et al. 2022).

### Docking of I3M and pOHB in occluded homology models

To find the best glucosinolate binding mode interacting with NPF2.9, NPF2.9Δ8#3 and NPF2.10 occluded models, and get structural insights about the role of the 8 key residues, we docked I3M and pOHB in the models using the Glide software (Friesner et al. 2006, Halgren et al. 2004) and the Standard Precision (SP) function, obtaining 20 poses per docking simulation. Docking in all three models yielded a variety of poses with similar binding modes. Among the best binding poses in all three models we identified conformations where I3M and pOHB are localized between the conserved arginine (R196 in NPF2.10, R156 in NPF2.9) and glutamate/aspartate (E513 in NPF2.10, D471 in NPF2.9), and all poses in proximity to the selectivity filter (Shown as a green line in Fig 10 and 11).

Binding poses in NPF2.10 show the sulfate group of both I3M and pOHB interacting with the positive charged arginine, and the sugar moiety interacts with the negative charged glutamate (Fig. 10 and 11). The side chains of I3M and pOHB are both close to the selectivity filter, but point in slightly different directions. Binding poses in NPF2.9 for I3M yielded similar poses as for NPF2.10, with the sulfate group interacting with the arginine, and the sugar moiety with the aspartate (Fig. 10). In contrast, the pOHB binding pose only showed the sulfate group interacting with the arginine while the sugar moiety points away from the aspartate (Fig. 11). This lack of interaction with the two conserved residues potentially explains the poor transport of pOHB by NPF2.9. Supporting of this, is the less negative docking score obtained for NPF2.9 with pOHB. The docking scores displayed in Fig. 11 expresses how well the docking fit; more negative indicates better binding. The docking scores for pOHB in NPF2.10 is −7.08 kcal/mol, where the score for the binding mode of pOHB in NPF2.9 is −4.63 kcal/mol, which suggests a worse fit of the glucosinolate in NPF2.9.

**Fig. 10.**
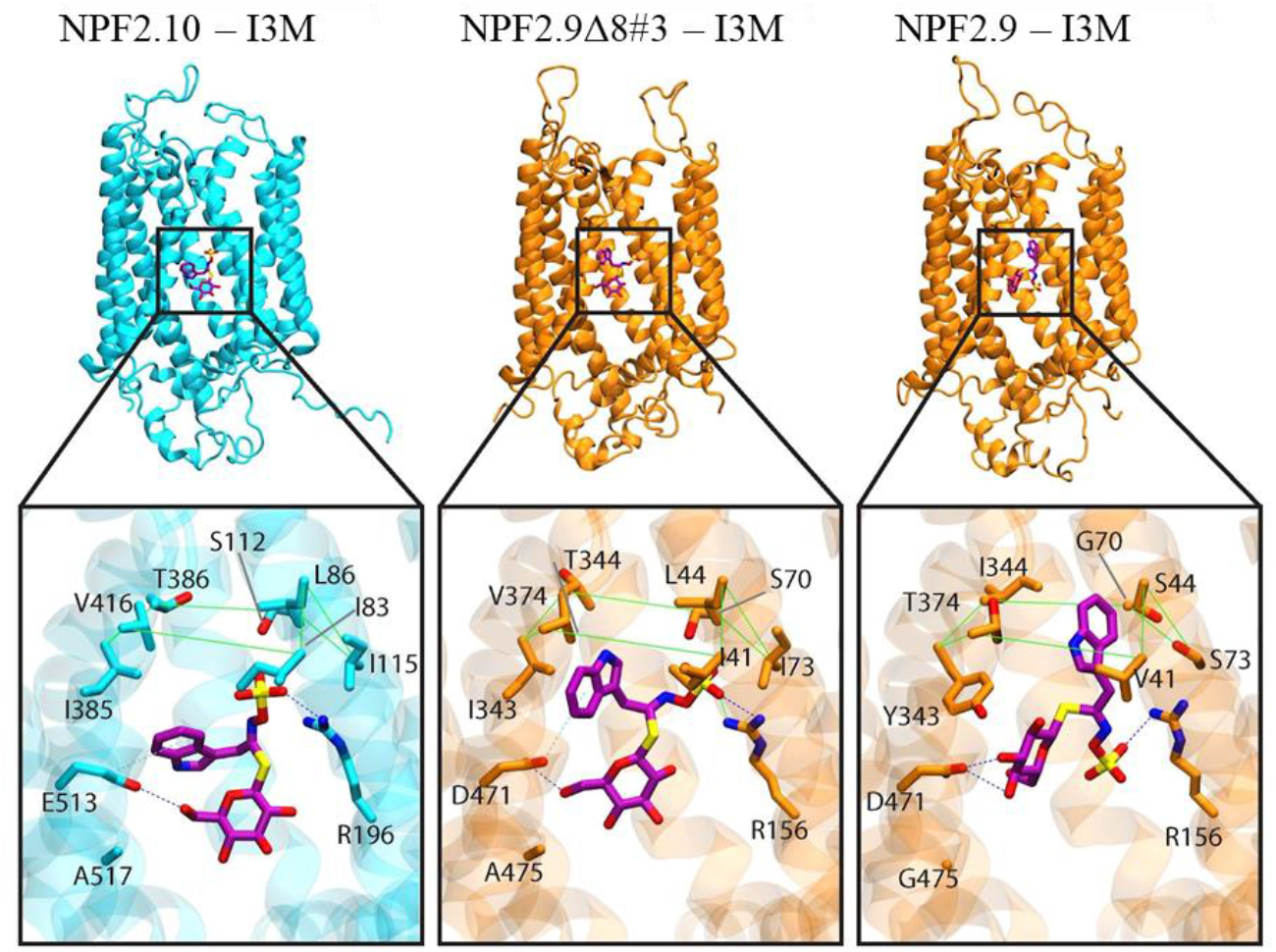
NPF2.10, NPF2.9Δ8#3 and NPF2.9 docking of I3M. NPF2.10 in cyan. NPF2.9 and NPF2.9Δ8#3 in orange. The selectivity “ring” is displayed as a green line. H-Bond are displayed as dotted blue lines, ionic interactions as dotted green lines, and aromatic H-Bonds as cyan dotted lines

The mutant NPF2.9Δ8#3 had a binding pose for I3M similar to both NPF2.9 and NPF2.10, with the sulfate group and sugar moiety interacting with the conserved residues. By visual inspection, the binding pose in the mutant is more similar to the binding pose observed for NPF2.10, than for NPF2.9, with the side chain pointing towards the left side of the green selectivity filter as for NPF2.10 (Fig. 10). The change in side chain position may lower the I3M binding capability of the transporter and fits with the transport of I3M being lower in the mutant than NPF2.9 (Fig. 8B). The binding pose of pOHB in the mutant homology model looks to be a more intermediate pose between the NPF2.9 and NPF2.10 binding poses (Fig. 11). The dockings scores achieved for the mutant with pOHB also suggest an intermediate binding, with a docking score of −6.32 kcal/mol, which is between the two wildtype transporters docking scores. This improved binding in the cavity of the mutant could explain why the introduction of the 8 NPF2.10 residues in the cavity of NPF2.9 can change the specificity of the transporter to be able to bind and transport pOHB.

**Fig. 11.**
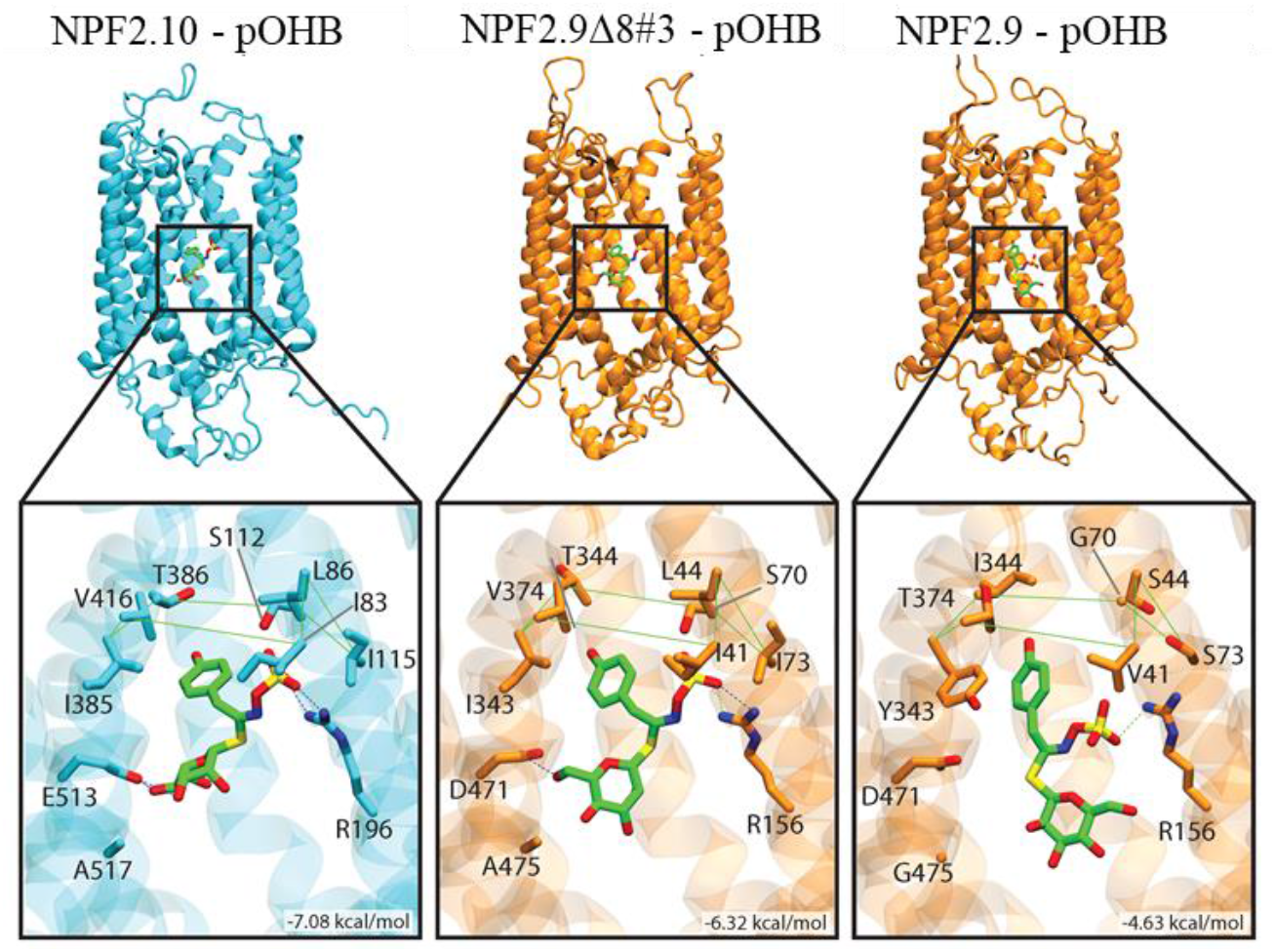
NPF2.10, NPF2.9Δ8#3 and NPF2.9 docking of pOHB. NPF2.10 in cyan. NPF2.9 and NPF2.9Δ8#3 in orange. The selectivity “ring” is displayed as a green line. H-Bond are displayed as dotted blue lines, ionic interactions as dotted green lines, and aromatic H-Bonds as cyan dotted lines

The docking score values should only be taken as indications, considering the general uncertainty associated with docking scores and that the dockings are based on homology models. Prediction of realistic binding energies requires high-resolution protein structures and post-processing of the poses by e.g. molecular dynamics simulations (Chen 2015, Guterres & Im 2020).

## Discussion

Substrate binding sites of transporters may in principle be inferred by introducing loss-of-function mutations. However, as also shown in this study, such results should be treated with caution, as a mutational approach may not only disrupt substrate binding, but may also disrupt the general transport mechanism or structural integrity of the protein. Gain-of-function mutations wherein substrate specificities are altered are less likely to suffer from misinterpretations, but they are also more rarely observed in the literature. Among major facilitator superfamily (MFS) transporters, a few cases have been successful. For example, the substrate specificity of a type II SUT was broadened to match that of type I SUTs through a gene shuffling method in combination with fluorescence-activated cell sorting (Reinders et al. 2012) and the D-xylose transporter Gal2 was made insensitive to inhibition by D-glucose through an error-prone PCR based cloning in combination with a growth based yeast screen (Farwick et al. 2014). In both cases, the successful outcome of the unguided mutational approach relied on methods for screening massive numbers of mutations. Examples of rational mutagenesis guided by homology models also exist. For example, docking the low efficient substrate L-arabinose into the cavities of Gal2 homology models built on different XylE structures in outward facing substrate bound conformations, revealed multiple gain-of-activity substitutions towards L-arabinose in a growth-based yeast screen (Wang et al. 2017).

Here, we succeeded in converting the narrow substrate preference of NPF2.9 towards that of NPF2.10 by mutating eight interdependent residues within the substrate binding cavity. We started out with 4.72*10^21^ possible mutation combinations (72 differentially conserved amino acid residues: 2^72^). By assessing their location in relation to potential substrate binding sites using two sets of homology models, we reduced the amount of mutant combinations to 4096 (2^12^). Based on a rational mutational approach wherein we first introduced all mutations simultaneously followed by individual reversions we identified the eight mutations by generating and screening only 30 mutants out of the 4096. This is less than 1 % of the mutants that could potentially confer the altered substrate dependent proton-coupling.

By introducing the mutations in NPF2.9, we selectively increased the preference for pOHB, while decreasing the preference for I3M, both to match the preferences seen in NPF2.10. The preference for 4MTB was increased as well, but only to an intermediate level compared to NPF2.10. The fact that we were also able to specifically reduce the pOHB transport of NPF2.10 through reciprocal mutations shows that the eight residues play a critical role in substrate recognition in the GTR-subclade in the NPF.

To investigate the structural changes in the mutant, we generated homology models of NPF2.9, NPF2.9Δ8#3 and NPF2.10 in an occluded state, and docked I3M and pOHB in these models. The dockings indicate strong interactions between the selectivity filter and the side chains for glucosinolates being transported and show that the core glucosinolate structure interacts with two conserved residues that are assumed to be involved in general substrate recognition in the POT family. Whereas interactions with the two conserved residues have previously been observed in dockings of I3M, 4MTB, 4MSB and benzyl in NPF2.10 using inward and outward facing conformations (Chung et al. 2022, Peña-Varas et al. 2022), but the selectivity filter has not been described before.

The overlapping yet distinct substrate specificities of the NPF2.9 and NPF2.10/2.11 have in this study proved to be a highly valuable model system for unraveling the mechanisms of substrate specificity. We have though our approach identified 8 residues important for substrate specificity in a NPF specialized metabolite transporter. The intermediate conversion of the 4MTB preference indicates that there are still more features determining substrate specificity in the GTR subclades to reveal. In this context, larger high-throughput mutational studies employing various synthetic fluorescent glucosinolates mimicking native substrate preferences (Manuscript I, this dissertation) represent a way forward.

Our findings lead us to propose a mechanistic model wherein the interaction of central features of potential substrates’ with conserved residues in the substrate binding cavity is determined by the interaction of the remaining and variable part of the substrates with a variable selectivity filter in the substrate binding cavity. Understanding why the specific composition of the selectivity filter favors specific glucosinolates and whether the selectivity filter represents a fundamental molecular basis for substrate recognition by the NPF family – potentially extending to the POT family, emerge as new burning questions. The knowledge gained will form an important prerequisite in the development of predictive tools for transporter- and substrate identifications and eventually lead to an ability to engineer specific substrates for specific transporters and vice versa.

## Materials and Methods

### Identifying differentially conserved amino acid residues

Homologues of *Arabidopsis thaliana* NPF2.9-NPF2.11 were identified by BLAST (Altschul et al. 1990) using the NCBI (Johnson et al. 2008) and Phytozome server v12 (Goodstein et al. 2012). The cutoff score was set to be the score of the closest of the other two *Arabidopsis thaliana* glucosinolate transporters (NPF2.9-NPF2.11). This led to retrieval of 39 glucosinolate transporter sequences; 13 NPF2.9 orthologues, 13 NPF2.10 orthologues and 13 NPF2.11 orthologues. An alignment was generated using Clustal Omega (Sievers et al. 2011), an UPGMA tree was generated using the Mega7 software (Kumar et al. 2016). 72 fully or partially fully differentially conserved amino acid residue positions were chosen aided by a Type II Divergence test (Gu & Vander Velden 2002).

### Identification of binding site residues (docking and HOLE analysis)

The NPF2.9 and NPF2.10 comparative models previously reported by our group (Jørgensen et al. 2017) were employed to study the pore-lining residues involved in the recognition and translocation of substrates in NPF2.9 and NPF2.10. We use two different approximations: *i*) Determination of residues facing NPF2.9 and NPF2.10 pores: To determinate the dimensions of the pore and to identify the pore-lining residues the algorithm HOLE (Smart et al. 1996) was employed in the VMD software (Humphrey et al. 1996) *ii*) Determination of potential residues involved in glucosinolate translocation: We docked 24 different glucosinolates (Supplementary Fig S1) into the NPF2.9 and NPF2.10 pore using Glide Standard Precision docking function (Friesner et al. 2006). The grid box was centered in the pore center (z axis) and expanded in both x- and y-axis to include the entire pore cavity as well as the residues that are facing the extra and intracellular space, which are possibly involved in recognition, specificity, and translocation mechanisms of the substrates. The final dimensions (Å) of the grid box were: Inner box (x,y,z): 10, 20, 35; Outer box (x,y,z): 30, 40, 55. The best 10 poses of each docked glucosinolate according the docking score, were selected for further analysis using the Cheminformatics Interaction Fingerprint tool in Maestro suite. Due to the only difference in the docked glucosinolates being the side-chain, in this analysis we selected the residues that interacted with this group, and the residues that interacted with other ligand moieties were scorned. Finally, both approximations (HOLE + Docking) results were combined and a set of residues (different between both NPF2.9 and NPF2.10) potentially involved in glucosinolate transport were proposed.

### Defining the sample mix-up control

Positon 70 of NPF2.9 was chosen and included as a 12^th^ position as a control for sample mix-up. It was identified as a central cavity residue (Wulff et al. 2019) and identified in the HOLE and docking analysis, but it is not stringently differentially conserved (Ser in all 26 NPF2.10 and NPF2.11 orthologous but among the 13 NPF2.9 sequences it is Ala one time, Gly eight times and Ser four times). Assuming that possibly hundreds of combinations of mutants would be tested and thus the workflow, including site-directed mutagenesis, DNA linearization, *in vitro* transcription, oocyte injection would be performed multiple times in large scale, mutants should function independently on the identity of position 70 being either Gly or Ser. That is, if two mutants only varying in the amino acid identity of position 70 do not display the same activity, we may assume sample mix-up.

Generation of occluded homology models of NPF2.10, NPF2.9 and NPF2.9Δ8#3

Alpha Fold model of AtNPF4.4 (Identifier AF-Q56XQ6-F1) was chosen as template for homology modeling of NPF2.9, NPF2.9Δ8#3 and NPF2.10 due to the occluded conformation of the predicted models. The models were created and validated using SWISS-MODEL (Benkert et al. 2011, Bertoni et al. 2017, Bienert et al. 2017, Guex et al. 2009, Mariani et al. 2013, Studer et al. 2014, 2020, 2021; Waterhouse et al. 2018). The NPF2.9, NPF2.9Δ8#3 or NPF2.10 protein sequences were used as target sequence, with AtNPF4.4 as template.

The models were relaxed using Molecular Dynamics as previously described (Peña-Varas et al. 2022) before docking. Briefly, all the models were optimized at pH 4.5 ± 2.0 with Epik (Shelley et al. 2007) and PROPKA (Olsson et al. 2011). Structures were minimized converging heavy atoms to RMSD = 0.3 Å, and then embedded into a pre-equilibrated phosphatidyl oleoylphosphatidylcholine (POPC) bilayer in a periodic boundary condition box with pre-equilibrated TIP3 water molecules. Na+ or Cl-ions were added to neutralize the systems, and then the ion concentration was set to 0.15 M NaCl. Prior to equilibrium simulations, the systems were relaxed using the default Desmond’s relaxation protocol. Then, the systems were equilibrated for 5 ns in an NPT ensemble at 300 K with the application of a restraint spring constant of 20 kcal·mol–1·Å–2 to the protein backbone atoms, followed with another 5 ns but reducing the restraint spring constant to 10 kcal·mol–1·Å–2. After a proper system equilibration, a 100 ns production MDs in NPT ensemble was performed applying a restraint spring constant to 1 kcal·mol–1·Å–2 to the secondary structure of each transporter. In both equilibrium and production MDs, temperature and pressure were kept constant at 300 K and 1.01325 bar respectively by coupling to a Nose–Hoover Chain thermostat (Cheng & Merz 1996) and Martyna–Tobias-Klein barostat (Martyna et al. 1994) with an integration time step of 2 fs. MDs were performed with Desmond (Bowers et al. 2006) and the OPLS3 force field (Harder et al. 2016). The simulations were analyzed with Desmond, KNIME, Schrödinger, and in-house scripts. Visualization was carried out with VMD (Humphrey et al. 1996) and Pymol (DeLano 2002).

### Docking of I3M and 4MTB in NPF2.10, NPF2.9 and NPF2.9Δ8#3 homology models

The ligands I3M and pOHB were constructed in Maestro and prepared by the LigPrep procedure (Sastry et al. 2013) prior to the docking. The Glide program (Friesner et al. 2006, Halgren et al. 2004) was used for docking of I3M and pOHB in the occluded models. The docking simultations were performed with the standard set-up in Glide using the SP scoring function. The substrate-binding site was defined by the residues lining the pore in the occluded conformations; in this way, we ensured that all residues that potentially interact with substrates were considered in the grid box, including the eighth key residues identified in this work. The molecular docking simulations were carried out with the otterbox edge of the grid setting as (32x × 32y × 42z) Å3. The best 20 docking solutions were considered for further analysis.

### Vector generation

GTR1/NPF2.10 (At3g47960), GTR2/NPF2.11 (At5g62680) and GTR3/NPF2.9 (At1g18880) in *Xenopus laevis* oocyte vector pNB1 (Nour-Eldin et al. 2006) were obtained from previously publications (Jørgensen et al. 2017, Nour-Eldin et al. 2012).

NPF2.9Δ12, NPF2.9Δ72, *Brassica rapa* NPF2.10 (Brara.A02314), *Brassica Rapa* NPF2.9 (Brara.F01308) *Capsella Rubella* NPF2.10 (Carub_v10016825mg) and *Capsella Rubella* NPF2.9 (Carub_v10012216mg) were obtained as codon optimized uracil containing non-clonal DNA fragments. NPF2.10Δ8 was obtained codon optimized already inserted into pNB1 as clonal gene from TWIST Bioscience.

All other mutations generated in this study were generated by USER fusion (Geu-Flores et al. 2007). Briefly, different fragments are generated though PCR. The first fragment in all combinations was amplified with a forward primer with the standard 5’ USER tail; the last fragment in all combinations was amplified with a reverse primer with the standard 3’ USER tail. All other primers used had an overlapping part with the next fragment. All PCR fragments had a uracil incorporated at both ends through the primers. Primers used for PCR amplification are listed in supplementary table S4.

### cRNA generation for oocyte injection

Linearized DNA templates for RNA synthesis were obtained by PCR amplifying the coding sequences surrounded by *Xenopus* 5’ – and 3’-UTRs from pNB1u using forward primer (5’ – AATTAACCCTCACTAAAGGGTTGTAATACGACTCACTATAGGG – 3’) and reverse primer (5’ − TTTTTTTTTTTTTTTTTTTTTTTTTTTTTATACTCAAGCTAGCCTCGAG – 3’). PCR products were purified using E.Z.N.A^®^ PCR Cycle Pure kit (OMEGA bio-tek) using the manufacturer’s instructions. Purified PCR products were *in vitro* transcribed with the mMessage mMachine T7 transcription kit (InVitrogen) using the manufacturer’s instructions. The yield of the transcription was measured on NanoDrop and diluted to a concentration of 500-550 ng/μL. Before use, the cRNA were run on gel electrophoresis.

### *Xenopus* oocyte influx transport assays

*Xenopus laevis* oocytes stage V-VI were purchased from Ecocyte Biosciences. They were injected with 50.6 nL cRNA (concentration 500-550 ng/μL) or nuclease free water using a Nanoinject II (Drummond Scientific Company). Glass capillars for the Nanoinject II were prepare with a needle puller and manually cut with surgical scissors, before they were filled with mineral oil. The injected oocytes were incubated 3 days at 16 °C in kulori pH 7.4 (90 mM NaCl, 1 mM KCl, 1 mM MgCl2, 1 mM CaCl2, 5 mM HEPES).

The injected oocytes were pre-incubated in kulori (90 mM NaCl, 1 mM KCl, 1 mM MgCl2, 1 mM CaCl_2_, 5 mM MES or HEPES) pH 4.5, 5, 5.5, 6, 7.4 or 8 as stated in figure legends, for 2 min, before being transferred to compound containing kulori of same pH for 1 hour. In the Export assay the oocytes was then furthermore in empty kulori of pH 5 or 7.4 for up to 3 hours.

After the assay the oocytes were washed three times in kulori pH 7.4 followed by one wash in Milli-Q water. Single oocytes were analysed; each oocyte were homogenized in 62.5 μL 50 % methanol containing the internal standard Sinigrin and stored minimum 1 hour at −20 °C. The homogenized oocyte samples were centrifuged at max speed for 10 min at 4 °C. 50 μL of the supernatant was mixed with 75 μL Milli-Q water to a final methanol concentration of 20 % before filtration through a 0.22 μm filter plate. The samples were analysed on LCMS.

### Quantification of glucosinolate content by LC-MS/MS

Oocytes samples were subjected to analysis by liquid chromatography coupled to tandem mass spectrometry. The method was modified from Crocoll et al. (2016) and parameters were adjusted and optimized to match the LC-MS/MS system in use (Crocoll et al. 2016). Briefly, chromatography was performed on a 1290 Infinity II UHPLC system (Agilent Technologies). Separation was achieved on a Kinetex XB-C18 column (100 x 2.1 mm, 1.7 μm, 100 Å, Phenomenex, Torrance, CA, USA). Formic acid (0.05%, v/v) in water and acetonitrile (supplied with 0.05% formic acid, v/v) were employed as mobile phases A and B respectively. The elution profile for glucosinolates was: 0-0.2 min, 5 % B; 0.2-3.5 min, 5-65 % B; 3.5-4.2 min 65-100 % B, 4.2-4.9 min 100 % B, 4.9-5.0 min, 100-5% B and 5.0-6.0 min 5 % B. The mobile phase flow rate was 400 μL/min. The column temperature was maintained at 40 °C. The liquid chromatography was coupled to an Ultivo Triplequadrupole mass spectrometer (Agilent Technologies) equipped with a Jetstream electrospray ion source (ESI) operated in negative ion mode. The instrument parameters were optimized by infusion experiments with pure standards. The ion spray voltage was set to 4500 V. Dry gas temperature was set to 325 °C and dry gas flow to 13 L/min. Sheath gas temperature was set to 400 °C and sheath gas flow to 12 L/min. Nebulizing gas was set to 55 psi. Nitrogen was used as dry gas, nebulizing gas and collision gas. Multiple reaction monitoring (MRM) was used to monitor precursor ion → fragment ion transitions. MRM transitions were determined by direct infusion experiments of reference standards. Detailed values for mass transitions can be found in supplemental Table S5. Both Q1 and Q3 quadrupoles were maintained at unit resolution. Mass Hunter Quantitation Analysis for QQQ software (Version 10, Agilent Technologies) was used for data processing. Linearity in ionization efficiency was verified by analyzing dilution series that were also used for quantification.

### Analysis of data

Data belonging to Fig. 6–9 were analysed using Rstudio. Concentrations from LCMS samples were calculated based on internal standard (Sinigrin) and response factor generated based on standard curves.

Response factor for oocytes was determined by standard curves for the compound investigated and the internal standard: response factor (f) = slope compound investigated/ slope internal standard.

Concentration in samples was calculated: (Area of peak for compound / Area of peak for internal standard) * 1/f * dilution factor * final internal standard concentration. Dilution factor: 156.25, final internal standard concentration: 0.5 μM. Outliers was removed automatically based on generated function in R that removes values outside of ±1.5*Inter quantile range. Statistical analysis is stated in figure legends. All were performed using ANOVA function aov() in R, followed by TUKEY post hoc test.

## Supplementary Material

### Supplementary figure

**Supplementary Fig. S1.**
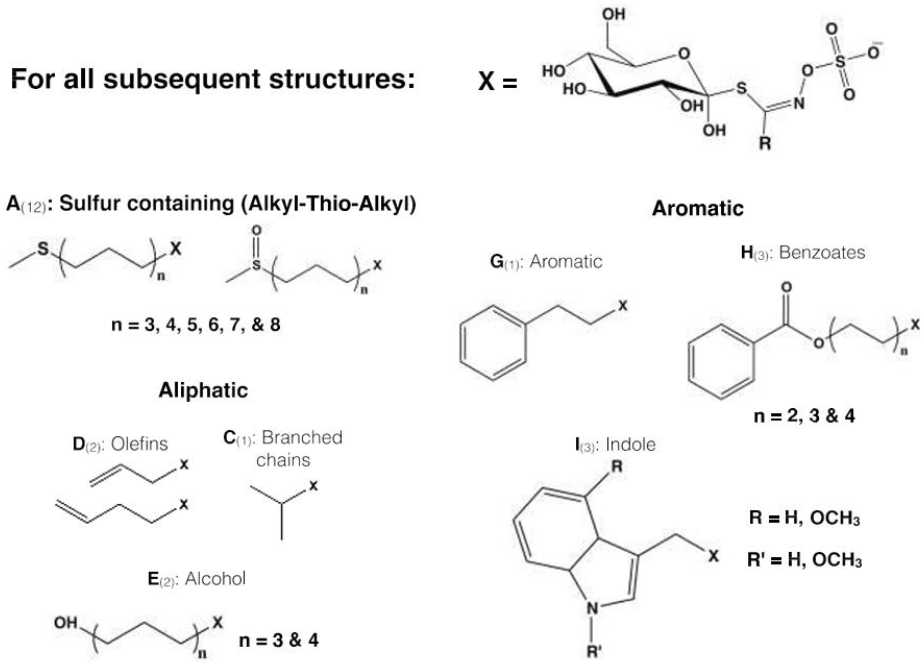
Glucosinolates docked into NPF2.9 and NPF2.10. A total of 24 glucosinolates: 12 sulfur containing aliphatic (A1 to A12), 5 sulfur deficient aliphatic (D1, D2, C1, E1, and E2), 7 Aromatic (G1, H1 to H3, I1 to I3). For I both R and R’ may not be OCH_3_ simultaneously.

**Supplementary Fig. S2.**
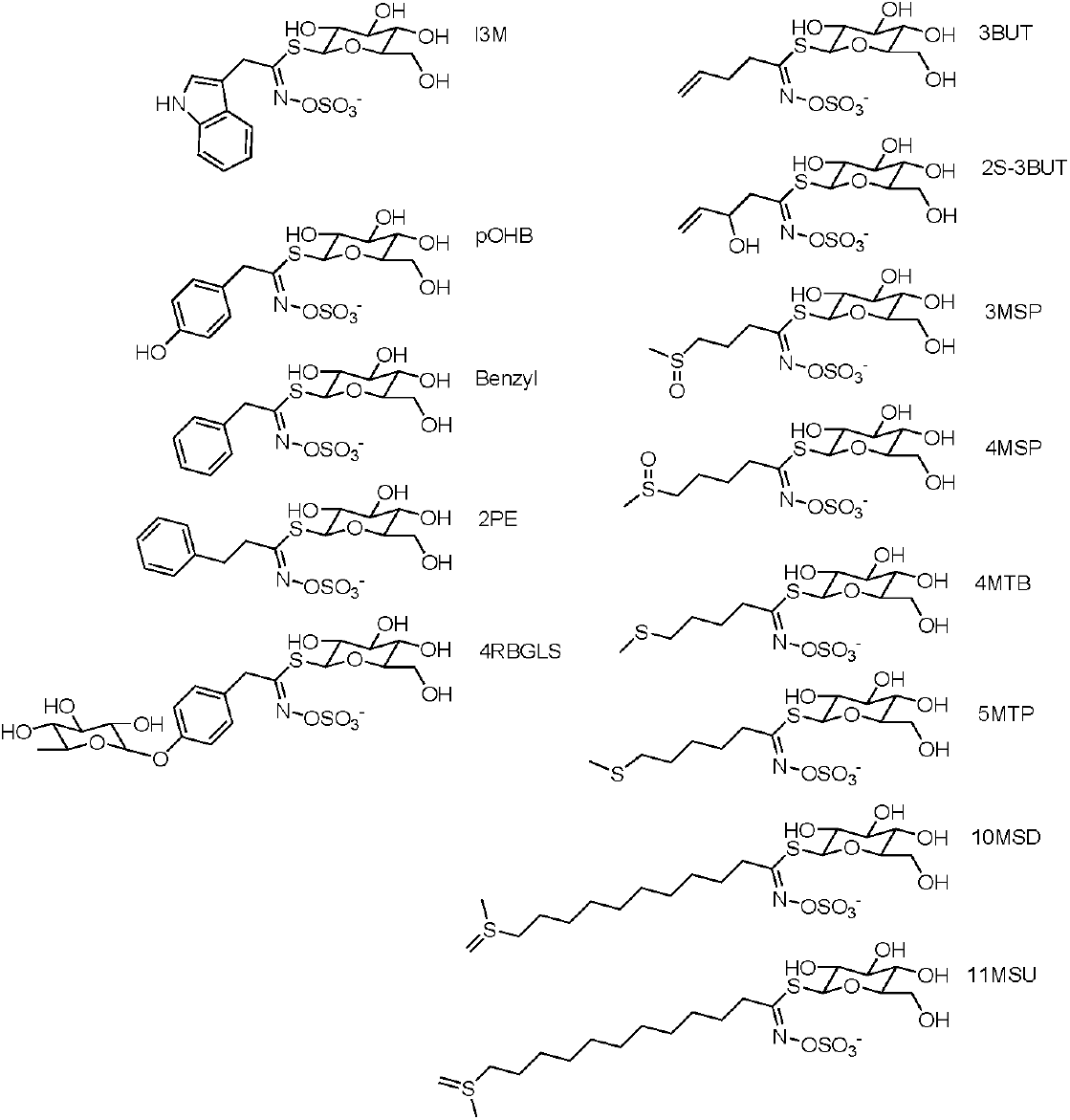
Glucosinolate structures of the 13 glucosinolates used in equimolar competition assay Fig 6.

### Supplementary Tables

**Supplementary table S1.**
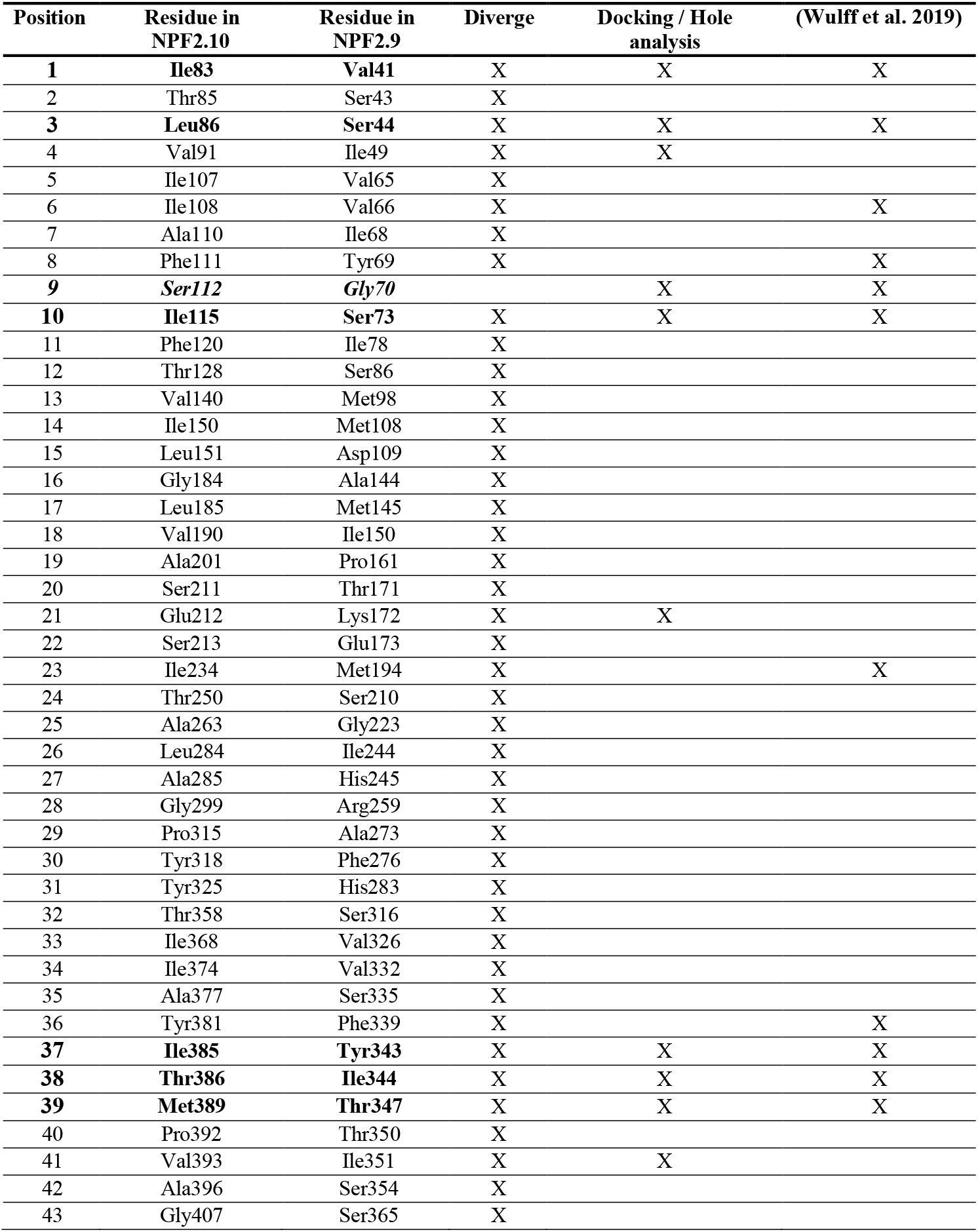
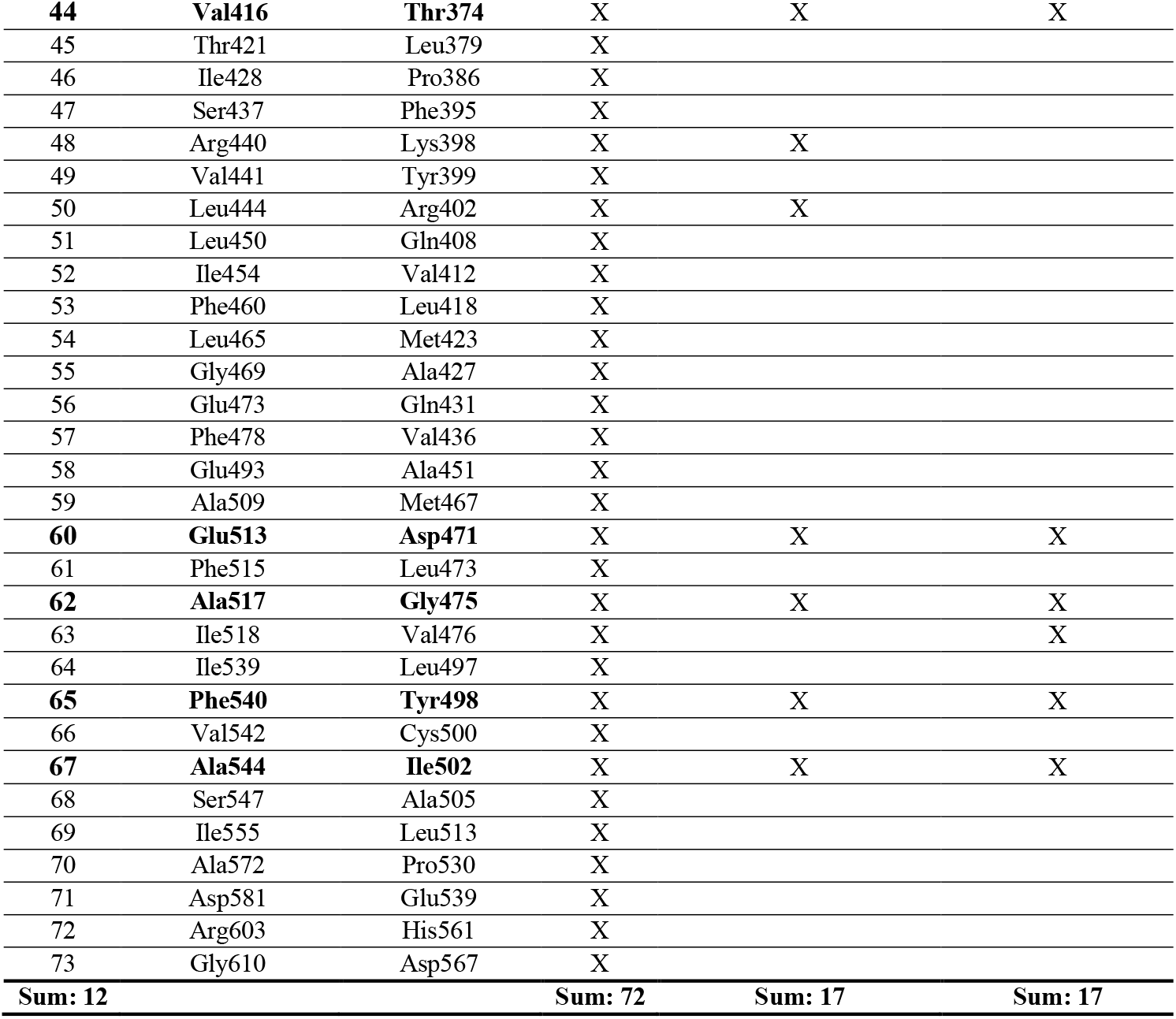
Amino acid positions. Residues marked in bold are the residues introduced in NPF2.9 to make up NPF2.9 Δ12. Position 9 written in italics is not particularly differentially conserved when comparing NPF2.9 to NPF2.10/NPF2.11, but was found in the two other models.

| Position | Residue in NPF2.10 | Residue in NPF2.9 | Diverge | Docking / Hole analysis | (Wulff et al. 2019) |
| --- | --- | --- | --- | --- | --- |
| **1** | **Ile83** | **Val41** | X | X | X |
| 2 | Thr85 | Ser43 | X |  |  |
| **3** | **Leu86** | **Ser44** | X | X | X |
| 4 | Val91 | Ile49 | X | X |  |
| 5 | Ile107 | Val65 | X |  |  |
| 6 | Ile108 | Val66 | X |  | X |
| 7 | Ala110 | Ile68 | X |  |  |
| 8 | Phe111 | Tyr69 | X |  | X |
| **9** | ***Ser112*** | ***Gly70*** |  | X | X |
| **10** | **Ile115** | **Ser73** | X | X | X |
| 11 | Phe120 | Ile78 | X |  |  |
| 12 | Thr128 | Ser86 | X |  |  |
| 13 | Val140 | Met98 | X |  |  |
| 14 | Ile150 | Met108 | X |  |  |
| 15 | Leu151 | Asp109 | X |  |  |
| 16 | Gly184 | Ala144 | X |  |  |
| 17 | Leu185 | Met145 | X |  |  |
| 18 | Val190 | Ile150 | X |  |  |
| 19 | Ala201 | Pro161 | X |  |  |
| 20 | Ser211 | Thr171 | X |  |  |
| 21 | Glu212 | Lys172 | X | X |  |
| 22 | Ser213 | Glu173 | X |  |  |
| 23 | Ile234 | Met194 | X |  | X |
| 24 | Thr250 | Ser210 | X |  |  |
| 25 | Ala263 | Gly223 | X |  |  |
| 26 | Leu284 | Ile244 | X |  |  |
| 27 | Ala285 | His245 | X |  |  |
| 28 | Gly299 | Arg259 | X |  |  |
| 29 | Pro315 | Ala273 | X |  |  |
| 30 | Tyr318 | Phe276 | X |  |  |
| 31 | Tyr325 | His283 | X |  |  |
| 32 | Thr358 | Ser316 | X |  |  |
| 33 | Ile368 | Val326 | X |  |  |
| 34 | Ile374 | Val332 | X |  |  |
| 35 | Ala377 | Ser335 | X |  |  |
| 36 | Tyr381 | Phe339 | X |  | X |
| **37** | **Ile385** | **Tyr343** | X | X | X |
| **38** | **Thr386** | **Ile344** | X | X | X |
| **39** | **Met389** | **Thr347** | X | X | X |
| 40 | Pro392 | Thr350 | X |  |  |
| 41 | Val393 | Ile351 | X | X |  |
| 42 | Ala396 | Ser354 | X |  |  |
| 43 | Gly407 | Ser365 | X |  |  |
| **44** | **Val416** | **Thr374** | X | X | X |
| 45 | Thr421 | Leu379 | X |  |  |
| 46 | Ile428 | Pro386 | X |  |  |
| 47 | Ser437 | Phe395 | X |  |  |
| 48 | Arg440 | Lys398 | X | X |  |
| 49 | Val441 | Tyr399 | X |  |  |
| 50 | Leu444 | Arg402 | X | X |  |
| 51 | Leu450 | Gln408 | X |  |  |
| 52 | Ile454 | Val412 | X |  |  |
| 53 | Phe460 | Leu418 | X |  |  |
| 54 | Leu465 | Met423 | X |  |  |
| 55 | Gly469 | Ala427 | X |  |  |
| 56 | Glu473 | Gln431 | X |  |  |
| 57 | Phe478 | Val436 | X |  |  |
| 58 | Glu493 | Ala451 | X |  |  |
| 59 | Ala509 | Met467 | X |  |  |
| **60** | **Glu513** | **Asp471** | X | X | X |
| 61 | Phe515 | Leu473 | X |  |  |
| **62** | **Ala517** | **Gly475** | X | X | X |
| 63 | Ile518 | Val476 | X |  | X |
| 64 | Ile539 | Leu497 | X |  |  |
| **65** | **Phe540** | **Tyr498** | X | X | X |
| 66 | Val542 | Cys500 | X |  |  |
| **67** | **Ala544** | **Ile502** | X | X | X |
| 68 | Ser547 | Ala505 | X |  |  |
| 69 | Ile555 | Leu513 | X |  |  |
| 70 | Ala572 | Pro530 | X |  |  |
| 71 | Asp581 | Glu539 | X |  |  |
| 72 | Arg603 | His561 | X |  |  |
| 73 | Gly610 | Asp567 | X |  |  |
| **Sum: 12** |  |  | **Sum: 72** | **Sum: 17** | **Sum: 17** |

**Supplementary table S2.** NPF2.9 Mutant overview. First letter corresponds to the identity of the residue in *Arabidopsis thaliana* NPF2.9. The number corresponds to the amino acid position in NPF2.9. The second letter corresponds to the identity of the residue in Arab*idopsis thaliana* NPF2.10. X marks in which mutants the given mutation is present.

| Mutant | V41I | S44L | G70S | S73I | Y343I | I344T | T347M | T374V | D471E | G475A | Y498F | I502A | Present in Fig |
| --- | --- | --- | --- | --- | --- | --- | --- | --- | --- | --- | --- | --- | --- |
| NPF2.9Δ12 | X | X | X | X | X | X | X | X | X | X | X | X | 4, 5AB |
| NPF2.9Δ11#1 |  | X | X | X | X | X | X | X | X | X | X | X | 5A |
| NPF2.9Δ11#2 | X |  | X | X | X | X | X | X | X | X | X | X | 5A |
| NPF2.9Δ11#3 | X | X |  | X | X | X | X | X | X | X | X | X | 5A |
| NPF2.9Δ11#4 | X | X | X |  | X | X | X | X | X | X | X | X | 5A |
| NPF2.9Δ11#5 | X | X | X | X |  | X | X | X | X | X | X | X | 5AB |
| NPF2.9Δ11#6 | X | X | X | X | X |  | X | X | X | X | X | X | 5A |
| NPF2.9Δ11#7 | X | X | X | X | X | X |  | X | X | X | X | X | 5AB |
| NPF2.9Δ11#8 | X | X | X | X | X | X | X |  | X | X | X | X | 5A |
| NPF2.9Δ11#9 | X | X | X | X | X | X | X | X |  | X | X | X | 5A |
| NPF2.9Δ11#10 | X | X | X | X | X | X | X | X | X |  | X | X | 5A |
| NPF2.9Δ11#11 | X | X | X | X | X | X | X | X | X | X |  | X | 5A |
| NPF2.9Δ11#12 | X | X | X | X | X | X | X | X | X | X | X |  | 5AB |
| NPF2.9Δ10#1 | X | X | X | X |  | X |  | X | X | X | X | X | 5B |
| NPF2.9Δ10#2 | X | X | X | X |  | X | X | X | X | X | X |  | 5B |
| NPF2.9Δ10#3 | X | X | X | X | X | X |  | X | X | X | X |  | 5B, 6, 7, 8AB |
| NPF2.9Δ9#1 | X | X | X | X |  | X |  | X | X | X | X |  | 5B, 7 |
| NPF2.9Δ9#2 |  | X | X | X | X | X |  | X | X | X | X |  | 7 |
| NPF2.9Δ9#3 | X |  | X | X | X | X |  | X | X | X | X |  |  |
| NPF2.9Δ9#4 | X | X |  | X | X | X |  | X | X | X | X |  | 7 |
| NPF2.9Δ9#5 | X | X | X |  | X | X |  | X | X | X | X |  | 7 |
| NPF2.9Δ9#6 | X | X | X | X | X |  |  | X | X | X | X |  | 7 |
| NPF2.9Δ9#7 | X | X | X | X | X | X |  |  | X | X | X |  | 7 |
| NPF2.9Δ9#8 | X | X | X | X | X | X |  | X |  | X | X |  | 7 |
| NPF2.9Δ9#9 | X | X | X | X | X | X |  | X | X |  | X |  | 7 |
| NPF2.9Δ9#10 | X | X | X | X | X | X |  | X | X | X |  |  | 7 |
| NPF2.9Δ8#1 | X | X | X | X | X | X |  |  |  | X | X |  | 8AB |
| NPF2.9Δ8#2 | X | X | X | X | X | X |  |  | X | X |  |  | 8AB |
| NPF2.9Δ8#3 | X | X | X | X | X | X |  | X |  | X |  |  | 8AB |
| NPF2.9Δ7#1 | X | X | X | X | X | X |  |  |  | X |  |  | 8AB |

**Supplementary table S3.** NPF2.10 Mutant overview. First letter corresponds to the identity of the residue in *Arabidopsis thaliana* NPF2.10. The number corresponds to the amino acid position in NPF2.10. The second letter corresponds to the identity of the residue in *Arabidopsis thaliana* NPF2.9. X marks in which mutants the given mutation is present.

| Mutant | I83V | L86S | S112G | I115S | I385Y | T386I | V416T | A517G | Present in Fig |
| --- | --- | --- | --- | --- | --- | --- | --- | --- | --- |
| NPF2.10Δ8 | X | X | X | X | X | X | X | X | 9 |

**Supplementary table S4.**
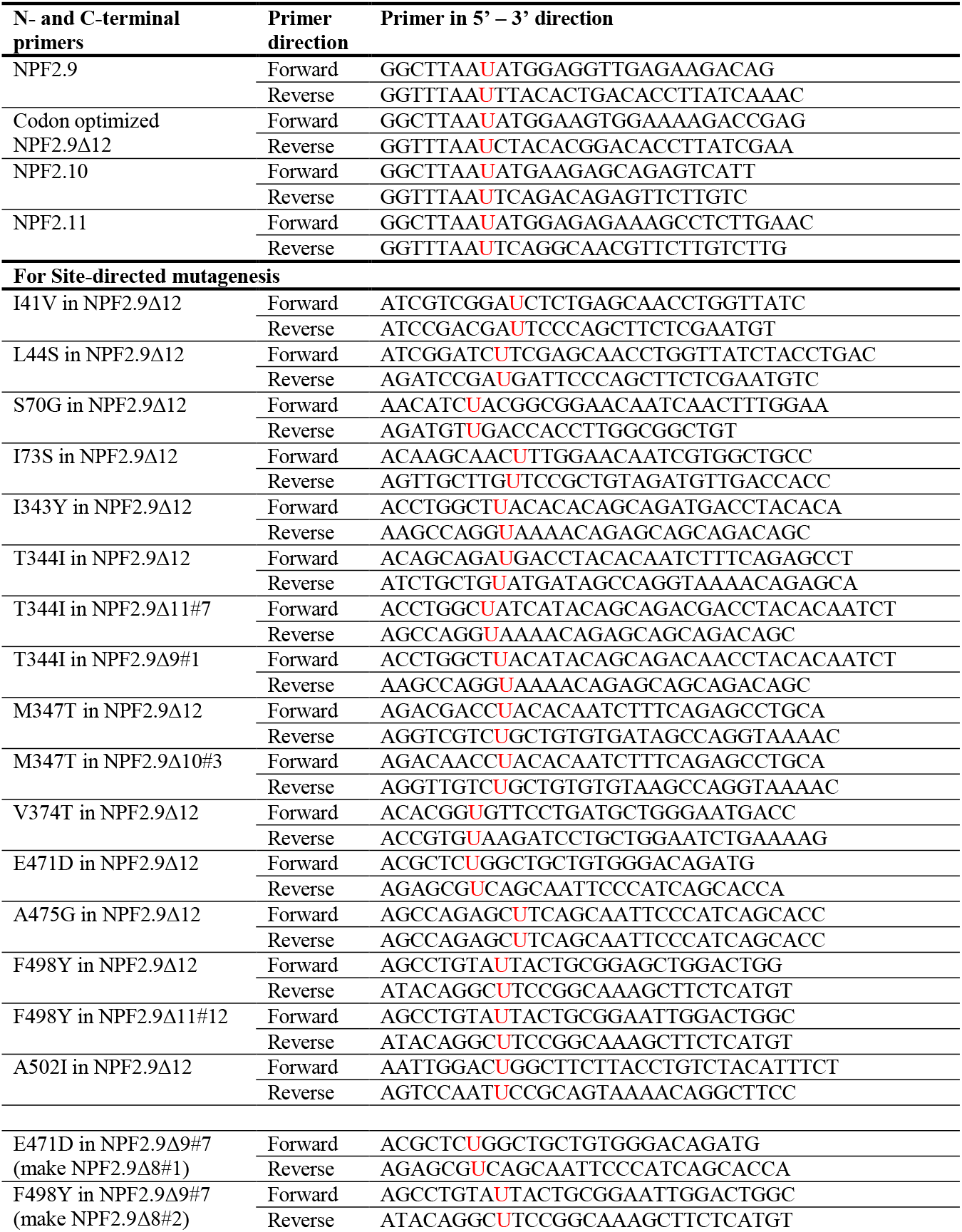
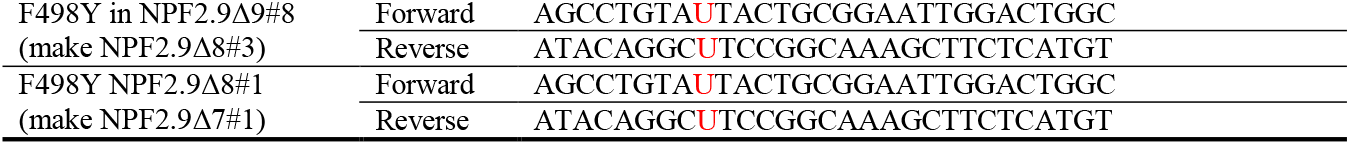
Primers used in this study. Uracils in primers marked with red.

**Supplementary table S5.** MRM transitions for glucosinolates quantified by LC-MS/MS. Qt = quantifier ion, additional transitions were used for identification.Q = quadrupole. CE = collision energy. IS = internal standard

| Analyte | Retention Time<br>[min] | Q1<br>[m/z] | Q3<br>[m/z] | Fragmentor<br>[V] | CE<br>[V] |
| --- | --- | --- | --- | --- | --- |
| 2-propenyl GLS (sinigrin) | 1.72 | 358.0 | 97.0<sup>Qt</sup> | 98 | 20 |
| [M-H]<sup>-</sup> |  | 358.0 | 259.0 | 98 | 16 |
| Internal standard (IS) |  | 358.0 | 75.0 | 98 | 36 |
| *p*-hydroxybenzyl GLS | 2.69 | 447.0 | 97.0<sup>Qt</sup> | 112 | 24 |
| (pOHB) |  | 447.0 | 259.0 | 112 | 24 |
| [M-H]<sup>-</sup> |  | 447.0 | 205.0 | 112 | 20 |
| 4mtb GLS | 2.56 | 420.0 | 97.0<sup>Qt</sup> | 107 | 24 |
| [M-H]<sup>-</sup> |  | 420.0 | 259.0 | 107 | 20 |
|  |  | 420.0 | 75.0 | 107 | 40 |
| I3M GLS | 2.69 | 447.0 | 97.0<sup>Qt</sup> | 112 | 24 |
| [M-H]<sup>-</sup> |  | 447.0 | 259.0 | 112 | 24 |
|  |  | 447.0 | 205.0 | 112 | 20 |

